# The purified recombinant FAT domain of the Focal Adhesion Kinase does not bind directly to recombinant Talin or MBD2 *in vitro*

**DOI:** 10.1101/2022.04.06.487260

**Authors:** Rayan Naser, Afaque A. Momin, Amal J. Ali, Jasmeen Merzaban, Stefan T. Arold

**Affiliations:** Bioscience Program, Biological and Environmental Science and Engineering Division, King Abdullah University of Science and Technology (KAUST), Thuwal 23955-6900, Kingdom of Saudi Arabia; Computational Biology Research Center, King Abdullah University of Science and Technology; Centre de Biologie Structurale (CBS), INSERM, CNRS, Université de Montpellier, F-34090 Montpellier, France

## Abstract

Controlled localization and activation of the focal adhesion kinase (FAK) functionally links adhesion, migration and survival of the cell. The C-terminal focal adhesion targeting (FAT) domain of FAK is an important regulator of the localization, activation and molecular associations of FAK. Here, we aimed to investigate the structural basis for how FAK FAT binds to Talin and MBD2, which were previously reported to be cytoplasmic and nuclear ligands, respectively. Using several biophysical methods with purified recombinantly expressed protein constructs, we failed to observe measurable interactions between FAT and either the Talin FERM domain or MBD2. We conclude that the association of FAT with these proteins requires additional factors or post-translational modifications not present in bacterially produced purified proteins.

## INTRODUCTION

Focal adhesions (FAs) are cellular adhesion sites that connect the cytoskeleton with the extracellular matrix (ECM) to maintain the multicellular structure. FAs convey mechanical changes in the extracellular environment into intracellular signaling pathways. FAs are large and dynamic multi-protein structures with transmembrane proteins, such as integrins, in addition to cytoskeletal and signaling proteins including vinculin, paxillin, talin, focal adhesion kinase (FAK) and others (1).

FAK is one of the key players at FAs. It is a non-receptor cytoplasmic protein tyrosine kinase (PTK) involved in the regulation of different cellular processes (mechanosensation, cell-cell signaling, cell polarization, migration, cell cycle and gene transcription) and in the maintenance of cellular structures (acto-myosin cytoskeletal dynamics, cell-cell junction formation and microtubule organization (2,3)). FAK expression and functions are precisely controlled by diverse mechanisms that include alternatively spliced forms, non-coding RNAs and an arsenal of interacting proteins. These factors directly or indirectly effect on the catalytic activity or protein stability of FAK (4–8). Given its various different functions and control mechanisms, overexpression or malfunction of FAK has been associated with many diseases (8). Consequently, understanding the molecular mechanisms coordinating the localization and activation of FAK may also open novel routes to therapeutically control FAK-driven cell metastasis and tissue invasion.

FAK is a 125kDa multi-domain protein that interacts with cell membrane receptors and transduces intracellular signals. It consists of an N-terminal band 4.1, ezrin, radixin, moesin (FERM) domain, a catalytic kinase domain and C-terminal focal adhesion targeting (FAT) domain. Long flexible linkers separate the central kinase domain from FERM (50 aa) and FAT (220 aa) domains **(Figure 1A)**. Although homologues of FERM, kinase and FAT domains are also present in other proteins, the specific combination and idiosyncrasies of these domains in FAK result in unique features (3,8).

**Figure 1:**
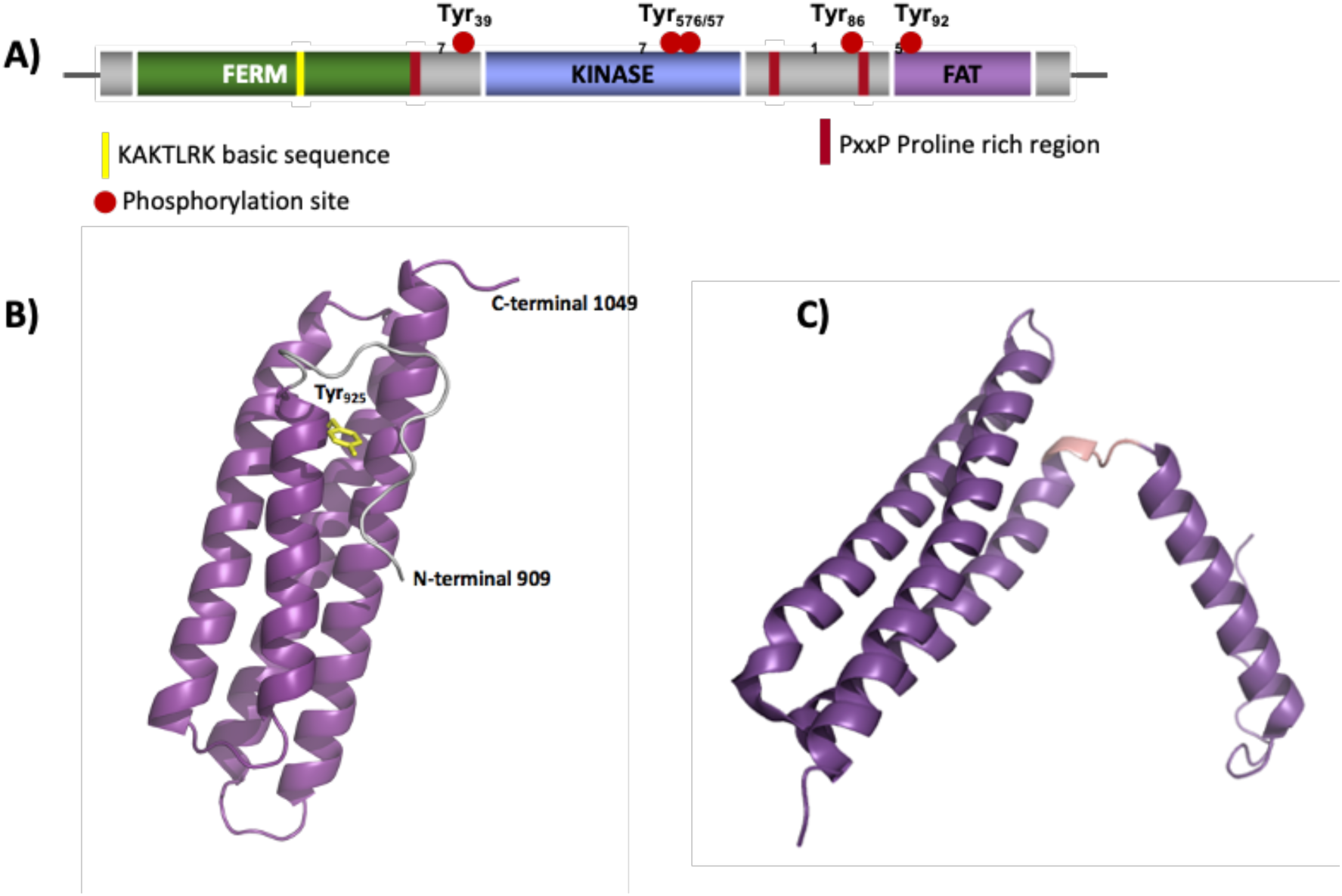
Overview of FAK domains and FAT structures. **A)** Schematic overview of FAK. The folded domains FERM, kinase and FAT are colored. Flexible linker regions are in gray. Major tyrosine phosphorylation sites are shown as red spheres. Proline rich regions are shown as red bars and the basic KAKTLRK patch is indicated by a yellow bar. **B)** The crystal structure of the FAT domain in its closed four-helix conformation (taken from PDB 3S9O). Helices 1 to 4 are represented in purple with the loops represented in gray. The N-terminal and C-terminal extensions are shown. The side chain of Y925 is shown in yellow. **C)** crystal form II of FAT domain intermediate with unfolded helix 1 separated from the helical bundle (PDB 1K04).

FAT comprises four amphipathic a-helices arranged into an antiparallel up-down-up-down 4-helix bundle **(Figure 1B)** (9,10). Helix 1 can dissociate from the rest of the domain, allowing phosphorylation of Tyr925 and subsequent binding of Grb2 via its SH2 domain **(Figure 1C)** (9). Thus, the open conformation is associated with an increased FA turnover by FAK (11). The circumstances that promote FAT Helix1 opening are not fully understood, but may involve ligand binding and bivalent ions (12).

FAT harbors docking surfaces for many ligands in different subcellular compartments and plays a critical role in the localization and regulation of the catalytic activity of FAK (3,13). Here, we aimed to investigate the structural and mechanistic basis for the interaction of FAT with two cellular ligands, Talin and MBD2.

Talin is a cytoskeletal adapter protein that functions at the cell–matrix attachment sites and binds to FAK. Although paxillin was first reported to be the protein that recruits FAK to the focal adhesion sites, transfected FAK was found to also co-immunoprecipitate with cellular Talin (14–16), and depletion of Talin disrupts the localization of FAK at FAs (16). These findings suggest that Talin plays a role in recruiting FAK mediating its phosphorylation and subsequent activation and localization. Conversely, Lawson et al. proposed that FAK is responsible for the recruitment of Talin to nascent adhesions (17). The exact binding model of these two proteins is not entirely understood. To further clarify the mechanism of the association between FAK and Talin, we investigated the interaction between FAK and Talin at the molecular level.

In addition to its roles at the cellular adhesions, FAK translocates into the nucleus where it binds to nuclear proteins and affects gene expression. By shuffling between the nucleus and the cell membrane, FAK can relay signals from the extracellular matrix and focal adhesions into the nucleus and affect gene regulation (18). Several physiological and chemical nuclear localization signals allow shuttling of FAK from the cytoplasm to the nucleus. Stress signals (such as oxidative stress), cell de-adhesion from the matrix and FAK inhibition are the main factors promoting FAK nuclear localization (19). FAK encompasses a lysine-rich nuclear localization signal (NLS) in the F2 lobe of the FERM domain and leucine-rich nuclear export signal (NES) in the kinase domain. FAK in its kinase inactive form exposes its NLS while the NES is masked by the FERM:Kinase interaction thus allowing FAK retention in the nucleus (18). Upon FERM-mediated transfer into the nucleus, FAK binds to p53 transcription factor and recruits the E3 ubiquitin-protein ligase Mdm2 promoting p53 degradation thus inhibiting p53-mediated gene regulation (20,21). In addition, nuclear FAK exerts anti-inflammatory effects by mediating the degradation of GATA4 and thus inhibiting the expression of vascular cell adhesion factor-1 (VCAM-1) (22). While binding to p53 and GATA4 is mediated via direct binding to the FERM domain of FAK, binding to another nuclear protein, the methyl CpG-binding domain protein 2 (MBD2), is mediated through FAT (23).

MBD2 binds methylated CpG-islands on the DNA and recruits HDAC1, thus regulating histone acetylation and chromatin structure (24,25). Binding of FAK to MBD2 is thought to reduce HDAC1 recruitment and, therefore, may influence regulation of chromatin structure and, subsequently, gene expression (23). The molecular basis of this interaction is not fully understood. Expression of different fragments of MBD2 in yeast, *in vitro* and in mammalian cells revealed that the GC-rich region and the methyl binding domain (MBD) are sufficient to bind to the FAT domain (23).

## 1. RESULTS

### 2. FAT constructs used

Unless noted differently, the recombinantly produced and purified FAT domain constructs used herein included FAK residues 892-1052. This FAT construct includes N-terminal residues preceding the four-helix bundle (residues 922 – 1049). These N-terminal residues stabilize the bundle structure through ionic bonds (R919 to D1039) and hydrophobic interactions (I909, P911, P912 and I917) (9,11). We have worked with this construct extensively in the past, and demonstrated its *in vitro* stability and capacity to bind ligands through crystallization, nuclear magnetic resonance (NMR) analyses, and biophysical binding studies (9,12,26).

### 3. Cloning of Talin F2F3

Talin is composed of a N-terminal head region (Talin-H1-433) and a highly elongated C-terminal rod domain (Talin-R434-2541) **(Figure 2A)**. Talin-H comprises a FERM domain consisting of three lobes, F1, F2, and F3 (27,28). Two studies have shown that the minimal binding site of FAK lies in the F2F3 domain of Talin (17,29), whereas other studies demonstrated that C-terminal 41 amino-acids of the FAT domain are sufficient for binding to Talin (10). Accordingly, Talin206-405, comprising F2 and F3 lobes of Talin, was cloned into m-pET32a. This Talin F2F3 construct had the C336S mutation to improve protein stability (30). Cell lysate was purified via Nickel affinity column chromatography and gel filtration **(Figure 2B)**. This construct displayed a high expression yield and a size exclusion chromatography elution profile corresponding to a 25 kDa protein **(Figure 2C)**. The protein was stable up to concentrations of 4 mg/ml.

**Figure 2:**
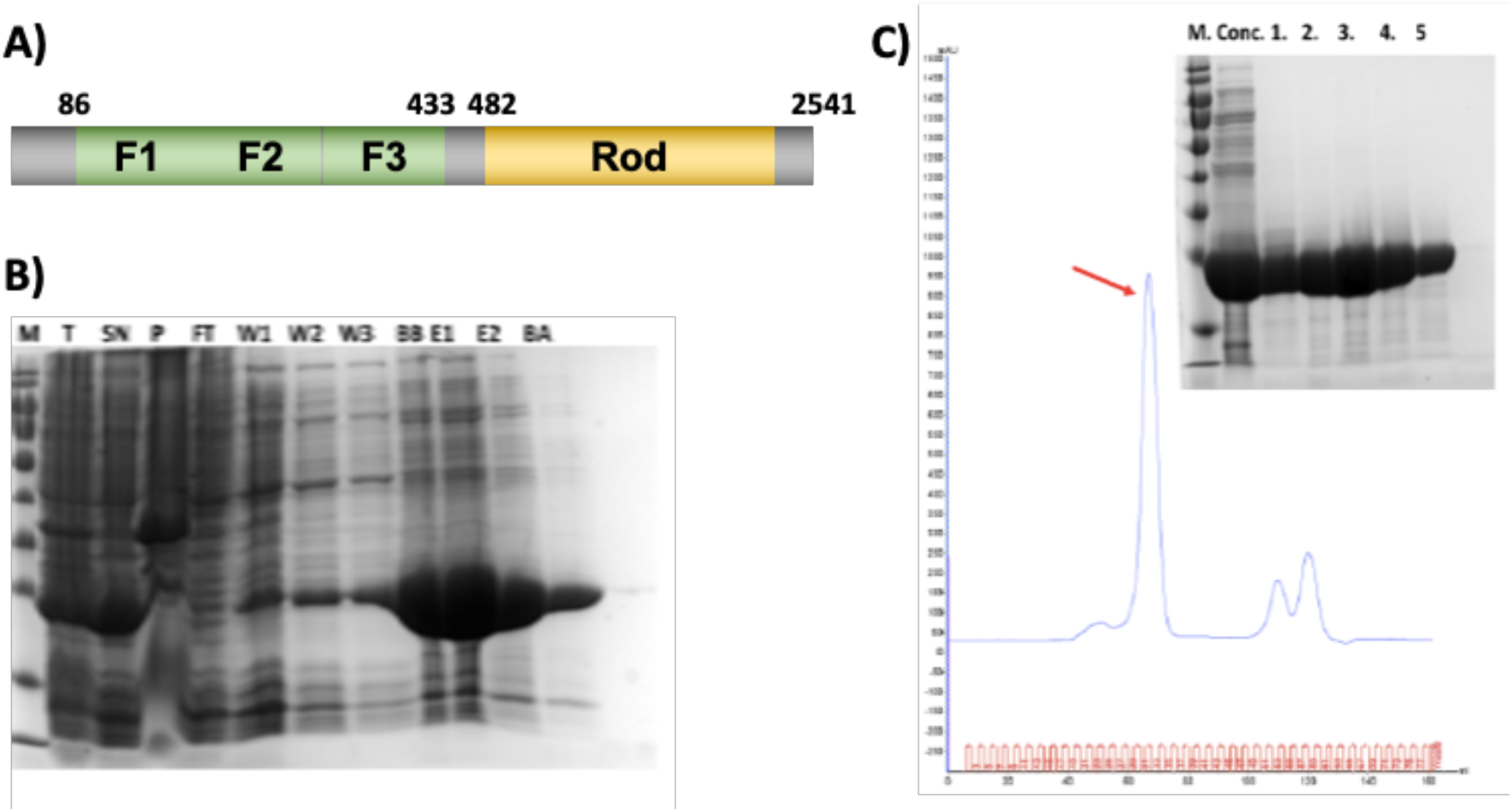
Purification of the Talin F2F3 fragment cloned into m-pET32a. **A)** Domain organization of Talin. Folded domains are colored, linker regions are in gray. **B)** SDS-PAGE gel showing the affinity purification of MBD2 fragment 3 (M: marker, T: total cell lysate, SN: supernatant, P: pellet, Ft: flow through of proteins not bound to Ni-beads, W1-W3: washes from 1 to 3 with wash buffer, BB: sample of beads before cleavage, E1-2: eluted protein fractions 1 and 2, BA: samples of beads after cleavage). **C)** Gel filtration chromatogram, red arrow showing the protein peak. *Inlay*: SDS-PAGE gel showing protein fractions collected after gel filtration (lanes 1 to 5 are fraction of the peak labelled with the red arrow).

We first tested the ability of his-tagged Talin F2F3 to pull down FAT in an assay in which Ni-beads and His-tagged Talin were incubated with FAT for 1 hour at room temperature. A control experiment was carried out without Talin where we expected to see no binding of FAT to Ni-beads. After several washes, Talin bound to Ni-beads failed to pull down FAT **(Figure 3A)**. We next performed a further analysis through immunoprecipitation assay. GST or Ni-Beads were washed with the corresponding buffer (CHAPS and Triton x-100) and incubated with both proteins overnight. We tested the ability of GST-FAT to retain His-Talin in the presence of GST beads **(Figure 3A)**, and the ability of His-Talin to retain GST-tagged FAT in the presence of Nickel beads **(Figure 3B)**. Proteins were stained with SYPRO Ruby for better sensitivity. Gel analysis revealed co-immunoprecipitation of GST-FAT with Talin bound to Ni-beads. We repeated the same experimental set-up and detected the precipitants by western blot. Using anti-His antibody, GST-FAT protein was not strongly efficient in retaining Talin from the supernatant **(Figure 3C)**. However, His-Talin was able to retain FAT from the supernatant using anti-GST antibody **(Figure 3C)**. To clarify the contradictory results between the simple, pull-down assay and co-immunoprecipitation and western blot, we tested whether GST-FAT would bind to Ni-beads even in the absence of Talin. Ni-beads (in triton x-100 buffer) and GST-FAT were incubated with and without His-Talin and immunoprecipitation was checked by western blot using anti-GST antibody. In both samples, with and without Talin, anti-GST antibody was able to detect GST-FAT immunoprecipitated with Ni-beads **(Figure 3D,E)**. Hence, GST-FAT bound non-specifically to Nickel beads.

**Figure 3:**
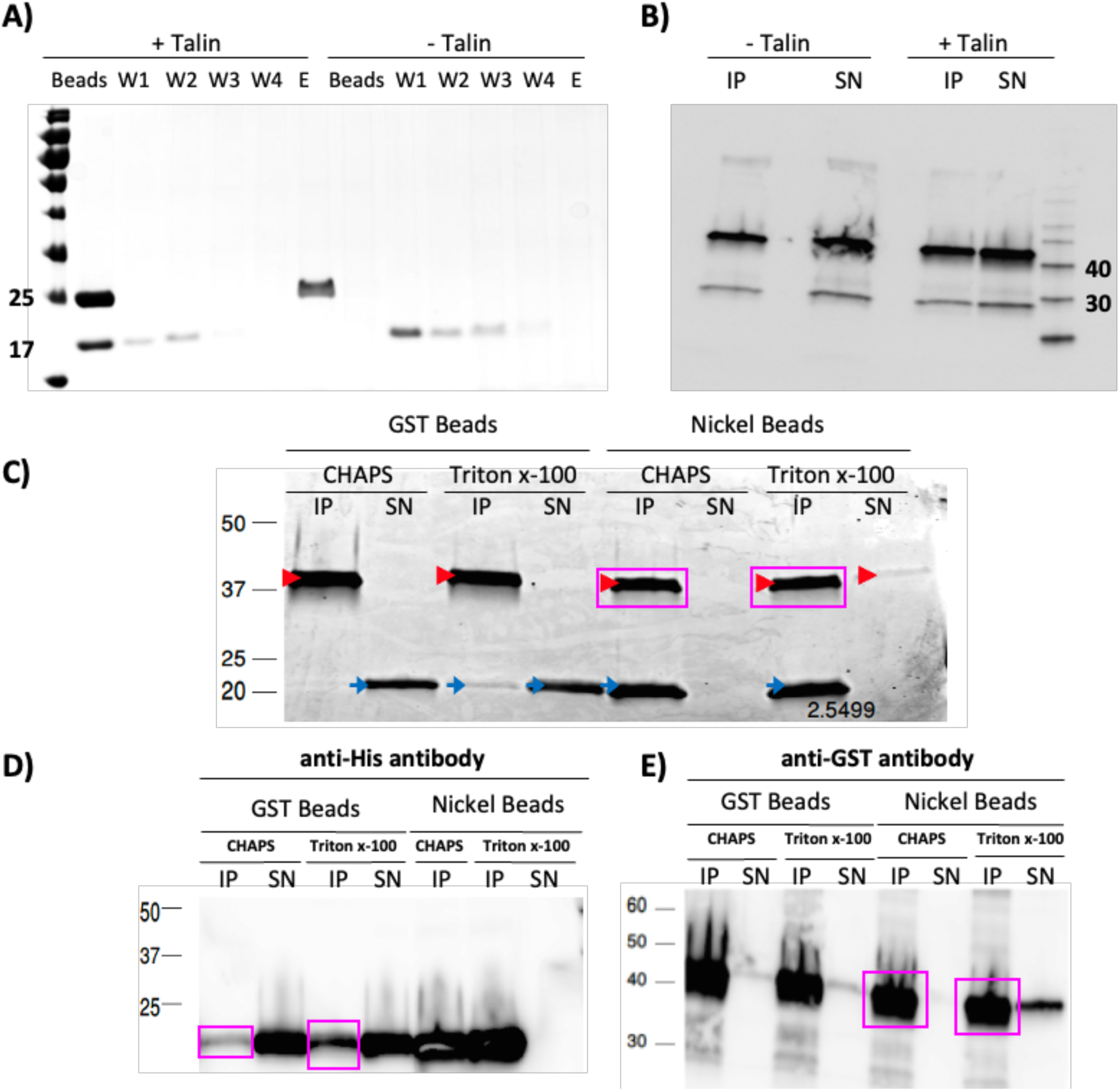
Pull-down assay failed to detect binding between FAT and Talin. **A)** The pull-down assay was carried out by incubating Nickel beads with either his-tagged Talin and FAT (+ Talin) or with FAT alone (-Talin). Beads were washed 4 times and then bound protein was eluted with 400mM imidazole. **B)** Immunoprecipitation on nickel beads of GST-FAT incubated with and without His-Talin. Detection by anti-GST antibodies revealed the unspecific binding of GST-FAT to Ni-beads (IP: immunoprecipitants, SN: supernatant). **C)** Immunoprecipitation: Beads were washed with the corresponding buffer and incubated with proteins overnight. GST-tagged FAT was used to retain His-tagged Talin in the presence of GST beads (left panel) and His-tagged Talin was used to retain GST-tagged FAT in the presence of Nickel beads (right panel). Proteins were stained with SYPRO Ruby. Red arrow is GST-FAT, blue arrow is His-Talin, and magenta boxes indicate GST-FAT immunoprecipitate with Talin. **D)** Western Blot using anti-His antibody shows that GST-FAT protein was not strongly efficient in retaining Talin (bordered with magenta box). **E)** Using anti-GST antibody, His-Talin incubated with Ni-beads was able to retain FAT from the supernatant (bordered with magenta box) (IP: immunoprecipitants, SN: supernatant).

#### Biophysical Analysis failed to show binding between FAT and Talin F2F3 in vitro

We next performed ITC titrations in which Talin and FAT samples were dialyzed in the same ITC buffer (20mM Phosphate buffer pH=7, 100mM NaCl, 2mM EDTA, 1mM TCEP). The titration of 500μM of FAT (placed in the syringe) onto 25μM of Talin (in the measurement cell) at 15C showed no variation in the peak size over the range of injections, which indicated that there was no binding under this condition **(Figure 4A)**.

**Figure 4:**
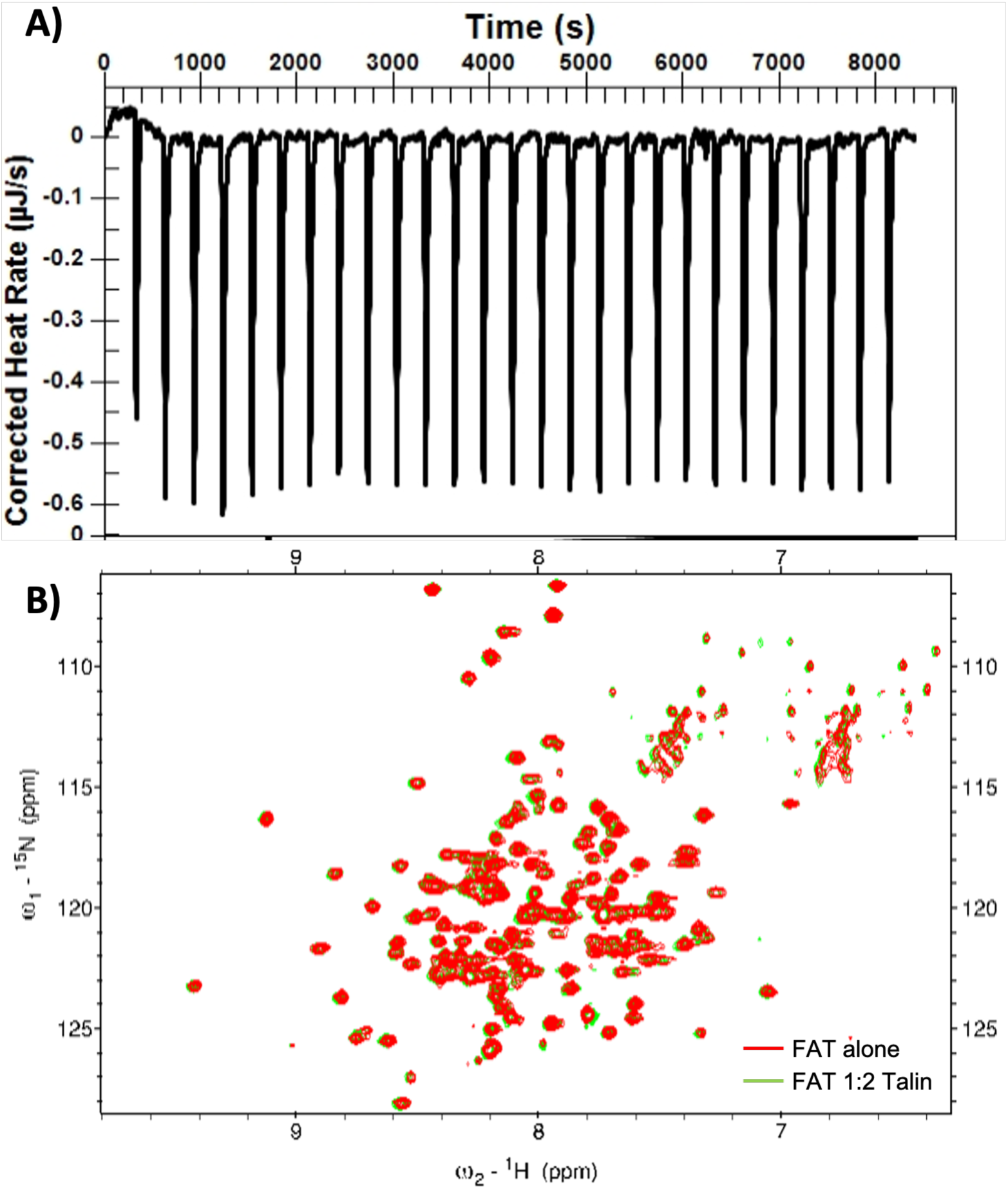
Biophysical analysis fails to show binding between FAT and Talin. **A)** Plot of ITC heats obtained over 27 injections of 400 μM FAT onto 25 μM Talin. No heat changes were observed during the titration, implying that no interactions took place. **B)** Overlay of ^1^H-^15^N HSQC experiments recorded on 15N-FAT alone (red) or in presence of a twofold excess of Talin (green). No significant chemical shift changes were observed.

Finally, we assessed the Talin-FAT interaction by NMR, through monitoring the chemical shift perturbations (CSPs) occurring when ^15^N-labelled FAT was incubated with Talin in ^1^H-^15^N HSQC experiments. 90 μM of ^15^N-labeled FAT were incubated with Talin at molar ratios of 1:0.5, 1:1 and 1:2. Consistent with the pull-down and ITC experiments, ^1^H-^15^N HSQC did not exhibit significant changes in the chemical shifts of FAT that was incubated with Talin when compared to apo-FAT (**Figure 4B**). We therefore concluded that under the conditions used, FAT and Talin did not interact, suggesting that their interaction, if it occurs, has a dissociation constants *Kd*, of more than 100 μM.

#### Cloning of the MBD2 fragments

To unveil the structural basis of the interaction between MBD2 and FAT, we designed MBD2 fragments of different lengths to test their binding affinities to FAT. MBD2 comprises a highly-disordered N-terminus with glycine-arginine (GR) repeats, a methyl binding domain (MBD) and a C-terminal coiled-coil (CC) region. We designed several MBD2 fragments to check the minimal binding site to FAT. MBD2 fragment 1 (MBD239-216) covers most of the GR-rich N-terminus and MBD, fragment 2 (MBD2121-216) contains a shorter GR-rich region along with MBD, whereas fragment 3 (MBD2146-216) constitutes solely the MBD **(Figure 5A)**.

**Figure 5:**
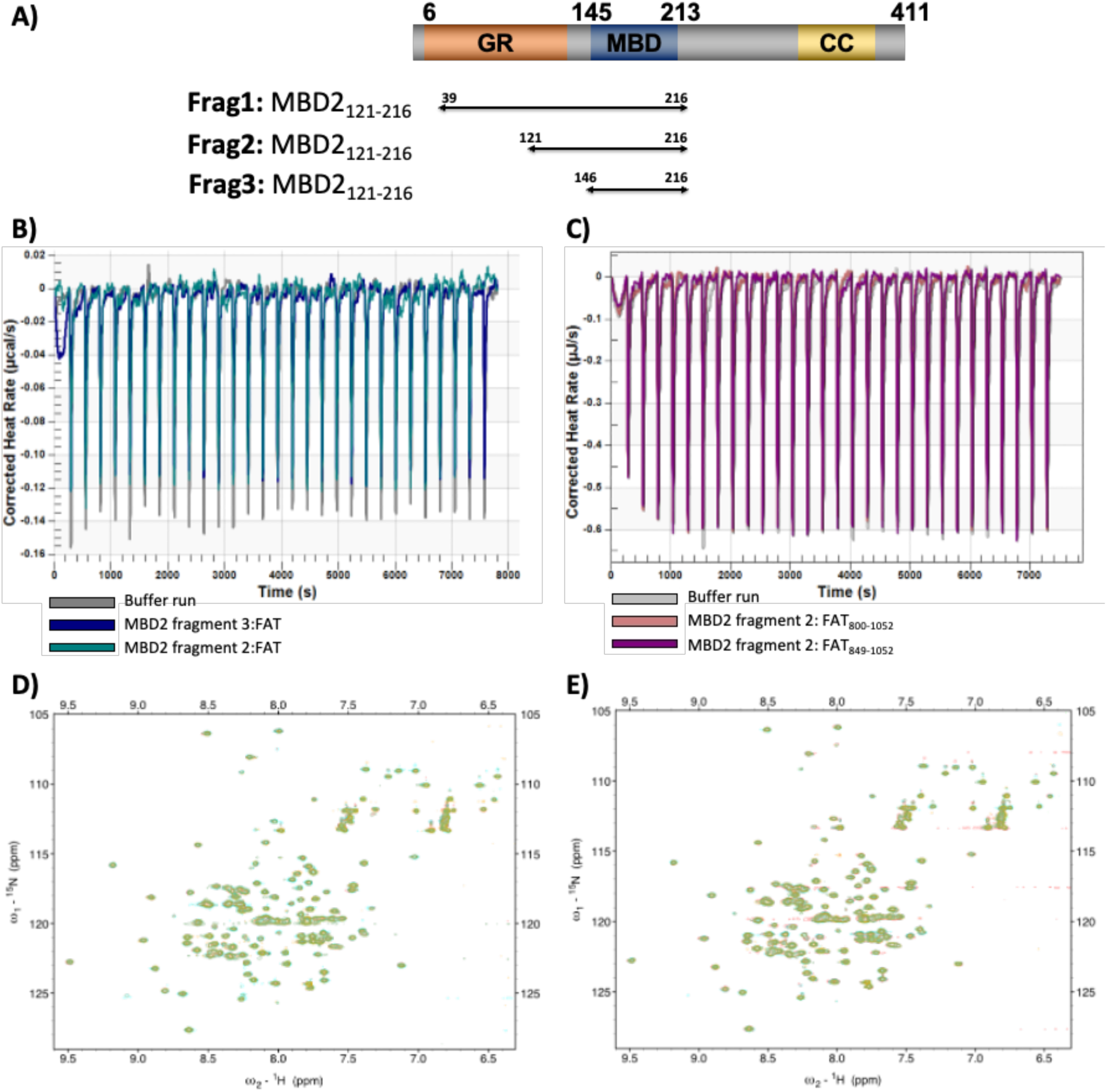
**A)** Design of MBD2 fragments **B**,**C) ITC titration experiments between MBD2 and FAT**. Shown are heats obtained during the injections of MBD2 onto FAT. **B)** ITC shows no binding between MBD2 fragment 2 and 3 with the FAT domain where 100 μM MBD2 were titrated onto 5 μM FAT. **C)** ITC shows no binding between MBD2 fragment 2 and two different FAT constructs where 50 μM MBD2 were titrated onto 5 μM FAT800 and FAT498. **D**,**E) NMR Titrations of FAT and MBD2:** Overlay of _1_H-_15_N-HSQC spectra of 100 μM _15_N-FAT in the absence (red) and presence of 0.2 (green), 1 (cyan) and 4 (yellow) times molar excess of MBD2 fragments. **D)** shows titrations of MBD2 fragment 3 and **E)** shows titrations of MBD2 fragment 2 into the FAT domain.

Initially, constructs were cloned into the pET28b vector with a His-tag. However, only fragment 2 was successfully cloned, expressed and purified with this vector. We then cloned the full-length protein into a pGEX6P1 vector with a GST-tag. The full-length clone was successfully transformed into BL21 cells, but expression tests failed. The following approach was to clone MBD2 (fragments and full length protein) into a pET32a vector as conducted by Desai et al. in (31). This approach proved more successful and we were able to clone MBD2 full length and shorter fragments **(Table 1)**. Of the four constructs cloned, only MBD2 fragment 2 and 3 showed good expression yields and were purified. After cell lysis and centrifugation, the lysate was incubated with Nickel beads. Elution was done by incubating cell lysate with 3C-protease to cleave MBD2 fragment from bound thioredoxin (Trx) and His-tags. For further purification, eluted protein was then applied to a gel filtration column.

**Table 1:**
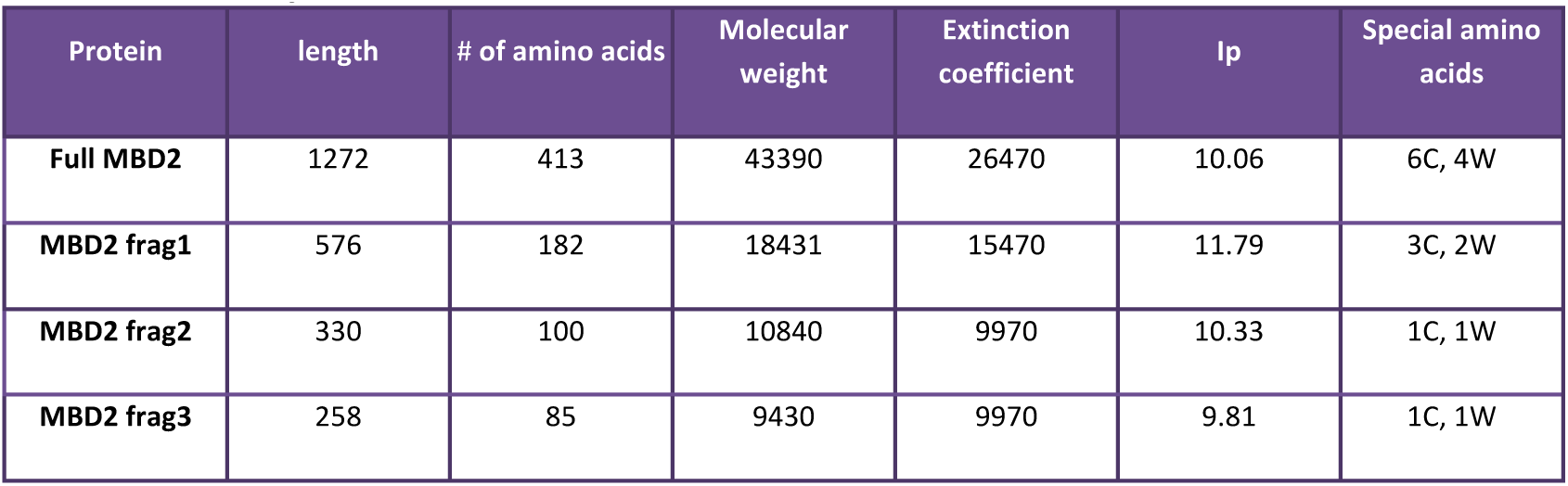
Molecular parameters of MBD2 constructs used

| Protein | length | # of amino acids | Molecular weight | Extinction coefficient | Ip | Special amino acids |
| --- | --- | --- | --- | --- | --- | --- |
| Full MBD2 | 1272 | 413 | 43390 | 26470 | 10.06 | 6C, 4W |
| MBD2 frag1 | 576 | 182 | 18431 | 15470 | 11.79 | 3C, 2W |
| MBD2 frag2 | 330 | 100 | 10840 | 9970 | 10.33 | 1C, 1W |
| MBD2 frag3 | 258 | 85 | 9430 | 9970 | 9.81 | 1C, 1W |

### 4. MBD does not bind to FAT892-1052 *in vitro*

We first performed ITC titrations in which MBD2 fragments 2 and 3 were dialyzed in the same ITC buffer (20 mM HEPES pH 7.5, 150 mM NaCl and 0.5 mM TCEP) with FAT892-1052. The titration of 100 μM MBD2 fragments onto 5 μM FAT at 15C showed no variation in the peak size over the range of peptide injections **(Figure 5B)**, which indicated that there was no binding with an affinity higher than ∼10-20 μM under our experimental conditions. To test if binding between FAT and MBD2 constructs occurred at lower affinities, we used NMR ^1^H-^15^N HSQC titration experiments as a robust and sensitive tool for testing low-affinity protein-protein interactions. However, the titration of MBD2 fragments 2 and 3 into 100 μM FAT showed no significant changes in the overlapped ^1^H-^15^N HSQC spectra **(Figure 5D,E)**.

Luo and his group have shown evidence of the FAT domain interaction with MBD2 both in yeast cells as well as *in vitro*. However, the shortest FAT construct used in their assay included the proline rich regions (PR) in the Kinase-FAT linker (23). Therefore we tested whether the longer FAT construct is essential for FAT:MBD2 interactions. Accordingly, FAT849-1052 containing PR2, and FAT800-1052 containing both PR1 and PR2 were cloned into a pJEx41 vector. However, ITC titrations of these longer FAT constructs with MBD2 fragment 2 did not show binding **(Figure 5C)**.

## 2. DISCUSSION

In this study, we aimed at deciphering the interaction between the FAT domain and two of its ligands, cytoplasmic Talin and nuclear MBD2. Both proteins were previously described as FAT binding partners; however, the structural basis of these interactions is not fully understood.

First, in *vitro* co-immunoprecipitation, ITC and HSQC titrations failed to detect an interaction between our recombinant FAT domain and the Talin F2F3 head. Although these findings were not consistent with FAT–Talin associations reported previously (28), *in vitro* binding experiments for FAK and Talin, in which both were recombinant proteins, were not performed previously. Rather, the interaction between FAK and Talin was detected through binding assays carried out in cell cultures, or involving proteins obtained both from cellular lysates, or from the transfection of one recombinant protein in cells that endogenously express the other protein (10,14–17,29). These studies have also tentatively mapped the FAT:Talin interaction. Initially, Chen et al. proposed the C-terminal residues 965 to 1012 of the FAT domain are sufficient for binding to Talin (14). Later, Lawson et al. showed that FAT E1015A mutation disrupts recruitment of Talin to nascent adhesions (17). Additionally, Talin225-357 has been described as the minimal FAK binding site on Talin, while a Talin construct containing additional residues, Talin186-357, binds to FAK with higher affinity (29).

Conversely, Cooley et al. were not able to detect an interaction between FAK and Talin using co-immunoprecipitation techniques for cellular proteins nor using *in vitro* binding assays with GST-fusion proteins (32). Interestingly, the only *in vitro* binding assay reporting an interaction between FAT and Talin involved *in vitro* translated FAK and GST-fusion Talin fragments (17). These observations may indicate that the interaction requires additional cellular partners. It is also possible that post-translational modifications are necessary for protein-protein interaction.

We were also unable to reproduce binding between the FAT domain and MBD2 fragments. Using pull-down and co-immunoprecipitation assays, Luo and his group provide evidence of FAK:MBD2 interactions through binding of the FAT domain of FAK to the MBD domain of MBD2. In this study, we used BL21-expressed recombinant proteins to study this interaction at the atomic level and for binding site mapping. *In vitro* biophysical binding assays showed that FAT892-1052 does neither bind to MBD2146-216 nor to a longer MBD2 construct containing a short GR-rich region along with MBD (MBD2121-216) under the experimental conditions used. These observations may be explained by two hypotheses. Firstly, the prokaryotic expression system used does not allow post-translational modifications needed to allow binding between the two domains. Indeed, Luo et al. used the mammalian expression system in their assays (23). In addition, it is possible that the domain boundaries used in our study do not harbor the minimal binding surface of interaction needed. Thus, additional FAK regions N-terminal to the ones we used might be required, and/or larger stretches of the intrinsically disordered region (IDR) of MBD2 which is connecting MBD to the CC domain of MBD2. IDRs can act as a center for various protein interactions (33), and the IDR of MBD2 is vital for the recruitment of histone deacetylase core components of the Nucleosome Remodeling and Deacetylase complex (NuRD) (31). Indeed, all MBD2 constructs used by Luo et al. encompassed the IRD.

In conclusion, we acknowledge that our work is insufficient to rule out a direct association between FAT and either Talin or MBD2. However, the negative results reported herein may facilitate designing future experiments to identify the nature of these associations.

## ACKNOWLEDGEMENTS

We thank the KAUST Bioscience, and Imaging and Characterization core labs for their assistance. This publication is based on the work supported by the King Abdullah University of Science and Technology (KAUST) Office of Sponsored Research (OSR) under the Award Number URF/1/2602-01-01.

## AUTHOR CONTRIBUTIONS

Protein cloning, expression and purification: RN. Biochemical and biophysical analysis: RN. NMR analysis: RN, AM. Cell biology, Western blots and pull-down: RN, AA, JM. Project design and supervision: STA, JM. Manuscript writing: RN, STA. All authors read and approved the manuscript.

## 2. MATERIALS AND METHODS

### 1. Cloning

The F2F3 lobe (206-405) of Talin FERM domain and full length MBD2 genes were codon optimized for bacterial expression and synthesized by Invitrogen (Thermo Fisher Scientific). Talin F2F3 domain had C336S mutation to improve protein stability and was cloned into a modified pET32a-m vector using the following oligonucleotides 5’-AAAACCTATGGTGTTAGCTTTTTTCTG-3’ and 3’-ATAACTGTAATAAGACTTTTTCTTTTTTAGTATT-5’ then transformed into *E. coli* BL21 (DE3). MBD2 Full length and smaller fragments were cloned into pET23a vector via Gibson cloning method using the following oligonucleotides: 5’-ATTGGATCCCTGGAAGTTCTGTTCCAGGGGCCCCTGCGTGCACATCCGGGTGGTGGT-3’ and 5’-ATTCTCGAGTTATCATGCTTCATCACCGCTATCCATTT-3’ for full length, 5’-ATTGGATCCCTGGAAGTTCTGTTCCAGGGGCCCCTGAGCGCACTGGCACCGAGTCCG-3’ as a

forward primer for fragment 1, 5’-ATTGGATCCCTGGAAGTTCTGTTCCAGGGGCCCCTGGCACCGCGTCGTGAACCGGTT-3’ a

forward primer for fragment 2, 5’-ATTGGATCCCTGGAAGTTCTGTTCCAGGGGCCCCTGGAAAGCGGTAAACGTATGGAT-3’ a forward primer for fragment 3 and 5’-ATTCTCGAGTTATTTGCTCGGCATCATTTTACCGGTAC-3’ as a reverse primer for the three fragments. Plasmids of MBD2 full length and smaller fragments were transformed into *E. coli* BL21 (DE3). His-tagged FAT (800-1052) was cloned by Twist Bioscience supplier in pJEx411c vector.

### 2. Protein expression and purification

FAT domain of human FAK (892-1052) was expressed as GST-fusion in *E. coli* BL21 using the expression vector pGex-6P2. Bacteria were grown in LB medium. FAT was expressed at 37°C for 3h while FAT (800-1052) was expressed at 30°C for 6h. MBD2 fragment 2 and 3 were expressed at 20°C overnight.

For purification, cells were thawed and lysed by mild sonication. Cell debris was removed by centrifugation. Lysates containing FAT were incubated with Glutathione Sepharose 4B beads (GE healthcare). The beads were thoroughly washed with 20 mM Tris-HCl (pH 8.0), 1 M NaCl, 2 mM EDTA, and 2 mM DTT. Beads were resuspended in 20 mM Tris-HCl (pH 8.0), 150 mM NaCl, 2 mM EDTA, and 2 mM DTT. Bound protein was incubated for 2 hr at room temperature with recombinant 3C protease.

Lysates containing His-tagged proteins were incubated with Ni-NTA agarose (invitogen). The beads were thoroughly washed with 20 mM HEPES (pH 7.5), 500 mM NaCl and 10mM imidazole. Beads were resuspended in 20 mM HEPES (pH 7.5) and 350mM NaCl and bound protein was incubated for 2 hr at room temperature with recombinant 3C protease. Eluted protein was dialyzed overnight with 20 mM HEPES (pH 7.5), 350mM NaCl and 2mM EDTA. Protein eluted from the affinity columns was further purified by size-exclusion chromatography using a Superdex 75 column (GE healthcare).

### 3. Isothermal titration calorimetry

To titrate FAT onto Talin, proteins were dialyzed in ITC buffer (20 mM Phosphate buffer pH 7, 100 mM NaCl, 2 mM EDTA, 1 mM TCEP). 1.5 ml of Talin were placed in the cell at a concentration of 25 μM. FAT at 400 μM concentration was placed in the injection syringe. For FAT:MBD2 titrations, proteins were dialyzed in 20 mM HEPES pH 7.5, 150 mM NaCl and 0.5 mM TCEP. 1.5 ml of FAT constructs were placed in the cell at concentrations of 5 μM. MBD2 fragments at 50 or 100 μM concentration were placed in 250 μl injection syringe.

Titrations were performed at 15°C with an initial injection of 2 μl, followed by 23 or 30 injections of 8 μl. ITC was performed on a Nano ITC from TA Instruments, and data were fitted using NanoAnalyze Software.

### 4. Pull-down assay

Simple Pull-down assay was carried out by incubating Nickel beads with either Talin and FAT or with FAT alone. Beads were directly washed 4 times with washing buffer (50mM HEPES pH=7.4, 150mM NaCl, 10% Glycerol and 1% triton-x) and bound protein was eluted with 400mM imidazole.

### 5. Immunoprecipitation and western blot

Beads were washed with the corresponding buffer and incubated with proteins overnight. GST-tagged FAT was used to fish out His-tagged Talin in the presence of GST beads and His-tagged Talin was used to fish out GST-tagged FAT in the presence of Nickel beads. Proteins were run on gels and stained with SYPRO Ruby or Invision His-tag or analyzed by western blot using anti-His antibody or anti-GST antibody.

### 6. Nuclear Magnetic Resonance

Cells were grown with ^15^N-NH4Cl dissolved in M9 minimal media solution, induced at OD=0.8 with 300uM IPTG and harvested after incubation overnight at 20°C (293.12 K). Protein samples were purified in 20mM HEPES, pH7.5, 150 mM NaCl, 1 mM TCEP buffer and ^15^N-labeled FAT protein at 100 μM was dissolved in a 10% D2O/90% H2O solution.

The samples were stable over the course of the NMR experiments. The 2D ^1^H-^15^N HSQC titrations (MBD2 and Talin) were performed at a temperature of 25 °C using a Bruker Avance III 700 MHz NMR spectrometer equipped with a triple resonance inverse TCI CryoProbe. Spectra were acquired with 2048 (^1^H) × 200-256 (^15^N) complex points, a spectral width of 16 ppm for ^1^H, 40 ppm for ^15^N, and averaged for 36-88 scans depending on sample concentration (80 to 100 *μ*M). ^1^H-^15^N HSQC spectra analysis was done using NMRFAM-SPARKY v1.4 software (Lee, Tonelli et al. 2015).

## REFERENCES

1. Zaidel-Bar R, Geiger B. The switchable integrin adhesome. J Cell Sci. 2010 May;123(Pt 9):1385–8.

2. Schaller MD. Cellular functions of FAK kinases: insight into molecular mechanisms and novel functions. J Cell Sci. 2010 Apr 1;123(Pt 7):1007–13.

3. Arold ST. How focal adhesion kinase achieves regulation by linking ligand binding, localization and action. Curr Opin Struct Biol. 2011 Dec;21(6):808–13.

4. Armendariz BG, Masdeu Mdel M, Soriano E, Urena JM, Burgaya F. The diverse roles and multiple forms of focal adhesion kinase in brain. Eur J Neurosci. 2014 Dec;40(11):3573–90.

5. Sulzmaier FJ, Jean C, Schlaepfer DD. FAK in cancer: mechanistic findings and clinical applications. Nat Rev Cancer. 2014 Sep;14:598–610.

6. Tai Y-L, Chen L-C, Shen T-L. Emerging roles of focal adhesion kinase in cancer. BioMed Res Int. 2015;2015:690690.

7. Panera N, Crudele A, Romito I, Gnani D, Alisi A. Focal Adhesion Kinase: Insight into Molecular Roles and Functions in Hepatocellular Carcinoma. Int J Mol Sci [Internet]. 2017 Jan 5;18(1). Available from: https://www.ncbi.nlm.nih.gov/pubmed/28067792

8. Naser R, Aldehaiman A, Diaz-Galicia E, Arold ST. Endogenous Control Mechanisms of FAK and PYK2 and Their Relevance to Cancer Development. Cancers Basel [Internet]. 2018 Jun 11;10(6). Available from: https://www.ncbi.nlm.nih.gov/pubmed/29891810

9. Arold ST, Hoellerer MK, Noble ME. The structural basis of localization and signaling by the focal adhesion targeting domain. Structure. 2002 Mar;10(3):319–27.

10. Hayashi I, Vuori K, Liddington RC. The focal adhesion targeting (FAT) region of focal adhesion kinase is a four-helix bundle that binds paxillin. Nat Struct Biol. 2002 Feb;9(2):101–6.

11. Kadare G, Gervasi N, Brami-Cherrier K, Blockus H, El Messari S, Arold ST, et al. Conformational dynamics of the focal adhesion targeting domain control specific functions of focal adhesion kinase in cells. J Biol Chem. 2015 Jan;290(1):478–91.

12. Hoellerer MK, Noble ME, Labesse G, Campbell ID, Werner JM, Arold ST. Molecular recognition of paxillin LD motifs by the focal adhesion targeting domain. Structure. 2003 Oct;11(10):1207–17.

13. Walkiewicz KW, Girault JA, Arold ST. How to awaken your nanomachines: Site-specific activation of focal adhesion kinases through ligand interactions. Prog Biophys Mol Biol. 2015 Oct;119(1):60–71.

14. Chen HC, Appeddu PA, Parsons JT, Hildebrand JD, Schaller MD, Guan JL. Interaction of focal adhesion kinase with cytoskeletal protein talin. J Biol Chem. 1995 Jul 14;270(28):16995–9.

15. Zheng C, Xing Z, Bian ZC, Guo C, Akbay A, Warner L, et al. Differential regulation of Pyk2 and focal adhesion kinase (FAK). The C-terminal domain of FAK confers response to cell adhesion. J Biol Chem. 1998 Jan 23;273(4):2384–9.

16. Wang P, Ballestrem C, Streuli CH. The C terminus of talin links integrins to cell cycle progression. J Cell Biol. 2011 Oct 31;195(3):499–513.

17. Lawson C, Lim ST, Uryu S, Chen XL, Calderwood DA, Schlaepfer DD. FAK promotes recruitment of talin to nascent adhesions to control cell motility. J Cell Biol. 2012 Jan 23;196(2):223–32.

18. Lim ST. Nuclear FAK: a new mode of gene regulation from cellular adhesions. Mol Cells. 2013 Jul;36(1):1–6.

19. Zhou J, Yi Q, Tang L. The roles of nuclear focal adhesion kinase (FAK) on Cancer: a focused review. J Exp Clin Cancer Res. 2019 Jun 11;38(1):250.

20. Golubovskaya VM, Finch R, Cance WG. Direct interaction of the N-terminal domain of focal adhesion kinase with the N-terminal transactivation domain of p53. J Biol Chem. 2005 Jul;280(26):25008–21.

21. Lim S-T, Chen XL, Lim Y, Hanson DA, Vo T-T, Howerton K, et al. Nuclear FAK promotes cell proliferation and survival through FERM-enhanced p53 degradation. Mol Cell. 2008 Jan 18;29(1):9–22.

22. Lim ST, Miller NL, Chen XL, Tancioni I, Walsh CT, Lawson C, et al. Nuclear-localized focal adhesion kinase regulates inflammatory VCAM-1 expression. J Cell Biol. 2012 Jun 25;197(7):907–19.

23. Luo SW, Zhang C, Zhang B, Kim CH, Qiu YZ, D. QS, et al. Regulation of heterochromatin remodelling and myogenin expression during muscle differentiation by FAK interaction with MBD2. EMBO J. 2009 Sep;28(17):2568–82.

24. Bird AP, Wolffe AP. Methylation-induced repression--belts, braces, and chromatin. Cell. 1999 Nov 24;99(5):451–4.

25. Leonhardt H, Cardoso MC. DNA methylation, nuclear structure, gene expression and cancer. J Cell Biochem Suppl. 2000;Suppl 35:78–83.

26. Alam T, Alazmi M, Naser R, Huser F, Momin AA, Astro V, et al. Proteome-level assessment of origin, prevalence and function of Leucine-Aspartic Acid (LD) motifs. Bioinformatics [Internet]. 2019 Oct 4; Available from: https://www.ncbi.nlm.nih.gov/pubmed/31584626

27. Rees DJ, Ades SE, Singer SJ, Hynes RO. Sequence and domain structure of talin. Nature. 1990 Oct 18;347(6294):685–9.

28. Garcia-Alvarez B, de Pereda JM, Calderwood DA, Ulmer TS, Critchley D, Campbell ID, et al. Structural determinants of integrin recognition by talin. Mol Cell. 2003 Jan;11(1):49–58.

29. Borowsky ML, Hynes RO. Layilin, a novel talin-binding transmembrane protein homologous with C-type lectins, is localized in membrane ruffles. J Cell Biol. 1998 Oct 19;143(2):429–42.

30. de Pereda JM, Wegener KL, Santelli E, Bate N, Ginsberg MH, Critchley DR, et al. Structural basis for phosphatidylinositol phosphate kinase type Igamma binding to talin at focal adhesions. J Biol Chem. 2005 Mar 4;280(9):8381–6.

31. Desai MA, Webb HD, Sinanan LM, Scarsdale JN, Walavalkar NM, Ginder GD, et al. An intrinsically disordered region of methyl-CpG binding domain protein 2 (MBD2) recruits the histone deacetylase core of the NuRD complex. Nucleic Acids Res. 2015 Mar 31;43(6):3100–13.

32. Cooley MA, Broome JM, Ohngemach C, Romer LH, Schaller MD. Paxillin binding is not the sole determinant of focal adhesion localization or dominant-negative activity of focal adhesion kinase/focal adhesion kinase-related nonkinase. Mol Biol Cell. 2000 Sep;11(9):3247–63.

33. Tompa P. Intrinsically disordered proteins: a 10-year recap. Trends Biochem Sci. 2012 Dec;37(12):509–16.

